## Supplementary Materials for "Structural basis for human mitochondrial tRNA maturation"

#### Supplementary Table 1. Cryo-EM data collection and refinement statistics

| Structure | **Complex 1** TRMT10C/SDR5C1 pre-tRNA$^{Ile}$ | **Complex 2** TRMT10C/SDR5C1 pre-tRNA$^{His\text{-}Ser}$ PRORP | **Complex 3** TRMT10C/SDR5C1 pre-tRNA$^{His(0,Ser)}$ ELAC2 | **Complex 4** TRMT10C/SDR5C1 pre-tRNA$^{Ile}$ TRNT1 |
|---|---|---|---|---|
| PDB code | | | | |
| EMDB code | | | | |
| **Data collection** | | | | |
| EM equipment | Titan Krios eBIC | FEI Titan Krios | FEI Titan Krios | FEI Titan Krios |
| Voltage (kV) | 300 | 300 | 300 | 300 |
| Detector | K3 | K3 | K3 | K3 |
| Mode | Counting, super resolution | Counting, super resolution | Counting super resolution | Counting, super resolution |
| Nominal magnification | 105,000x | 130,000x | 130,000x | 130,000x |
| Pixel size (Å per pixel area) | 0.723 | 0.652 | 0.652 | 0.652 |
| Electron dose, total (e-/ Å$^2$) | 40.10 | 48.52 | 48.90 | 42,20 |
| Defocus range (μm) | -0.9 to –2.5 | -0.9 to -2.4 | -0.9 to -2.4 | -0.8 to -2.2 |
| Exposure (s) | 3.0 | 1.34 | 1.34 | 1.24 |
| Frames | 40 | 150 | 50 | 46 |
| Number of micrographs | 6,474 | 9,826 | 23,314 | 10,835 |
| **Processing** | | | | |
| Software | Relion 3.1.0 | WARP CryoSparc v3.1 | WARP Cryosparc v3.1 | Relion v.3..1.0 CryoSparc v4.0 |
| Initial number of particles | 1,536,128 | 1,789,726 | 2,049,642 | 2,420,733 |
| Final number of particles | 141,653 | 81,396 | 74,166 | 25,554 |
| Symmetry imposed | C1 | C1 | C1 | C1 |
| Map resolution, $FSC_{0.143}$ (Å) | 3.20 | 2.76 | 2.79 | 3.14 |
| Map resolution range (Å) | 2.6-6.5 | 2.4-5.9 | 2.4-6.2 | 2.4-6.2 |
| Map-sharpening B-factor (Å$^2$) | -133.7 | -70.7 (Map 1) -246.1 (Map 3) | -69.9 (Map 1) -191.3 (Map 3) | -60.7 (Map 1) -183.4 (Map 2) |
| **Model composition** | | | | |
| Non-hydrogen atoms | 10626 | 16223 | 17251 | 14576 |
| Protein residues | 1232 | 1834 | 2007 | 1701 |
| RNA bases | 58 | 93 | 86 | 66 |
| Ligand | 5 | 7 | 6 | 6 |
| **Refinement** | | | | |
| Initial model used (PDB code) | 7ONU, 5NFJ, Complex 4 | 7ONU, 5NFJ AF-Q7L0Y3, AF-O15091, Complex 3 | 7ONU, 5NFJ, AF-Q7L0Y3, AF-Q9BQ52, Complex 2 | 7ONU, 5NFJ, AF-Q7L0Y3, AF-Q96Q11, Complex 1 |
| Software | Phenix-1.20.1-4487 | Phenix-1.20.1-4487 | Phenix-1.20.1-4487 | Phenix-1.20.1-4487 |
| Correlation coefficient, masked | 0.81 | 0.84 | 0.75 | 0.83 |
| **Model validation** | | | | |
| MolProbity score | 1.58 | 1.61 | 1.57 | 1,66 |
| All-atom clash score | 5.37 | 6.17 | 5.97 | 7.65 |
| Poor rotamers | 0.00 | 0.00 | 0.00 | 0.00 |

| | | | | |
|---|---|---|---|---|
| *Ramachandran statistics (%):* | | | | |
| Favoured (overall) | 95.74 | 95.99 | 96.34 | 96.38 |
| Allowed (overall) | 4.26 | 4.01 | 3.66 | 3.62 |
| Outliers (overall) | 0.00 | 0.00 | 0.00 | 0.00 |
| *RMS deviations:* | | | | |
| Bond length (Å) | 0.003 | 0.002 | 0.002 | 0.003 |
| Bond angle (°) | 0.480 | 0.476 | 0.486 | 0.446 |

### Supplementary Figures

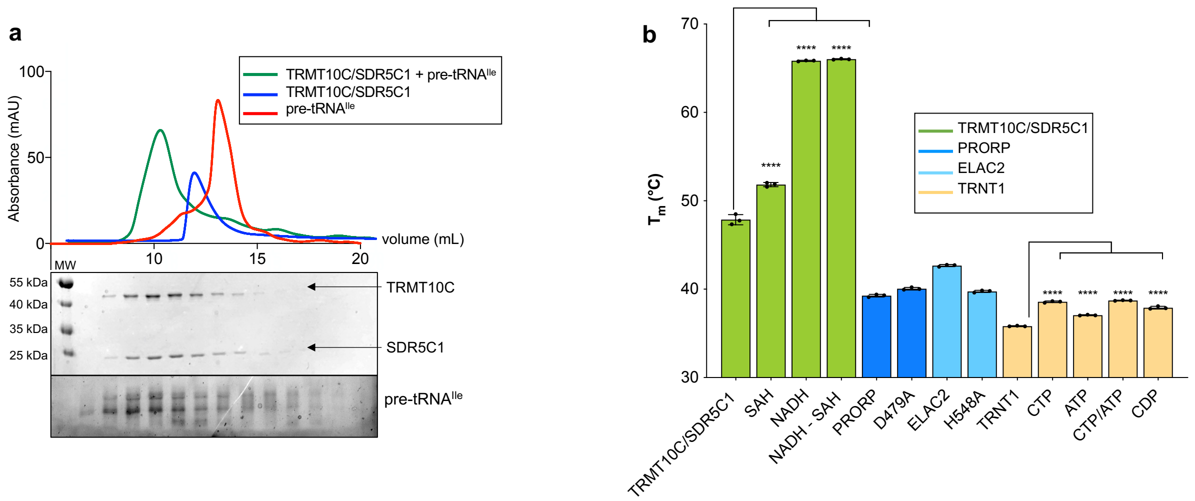

**Supplementary Fig. 1: Biochemical characterization of TRMT10C/SDR5C1/tRNA$^{Ile}$ complex and stability of the maturation enzymes in presence of their cofactors or analogues. a** Analytical size exclusion chromatography (Superdex 200 increase 10/300 GL) of TRMT10C/SDR5C1 in presence of pre-tRNA$^{Ile}$ which confirms that TRMT10C/SDR5C1 forms a stable complex with pre-tRNA$^{Ile}$, 14% SDS-PAGE gel analysis of the fractions. **b** Melting temperatures (Tm, in °C) measured from DSF experiments of TRMT10C/SDR5C1 (in green), PRORP and $PRORP_{D479A}$ (in dark blue), ELAC2 and $ELAC2_{H548A}$ (in cyan) and TRNT1 (in beige) with no ligand (Apo) or in the presence of cofactors or analogues. P-values are indicated as follows: ns = P > 0.05, * = P 0.05, *** = P 0.001 and **** = P 0.0001, for n = 3.

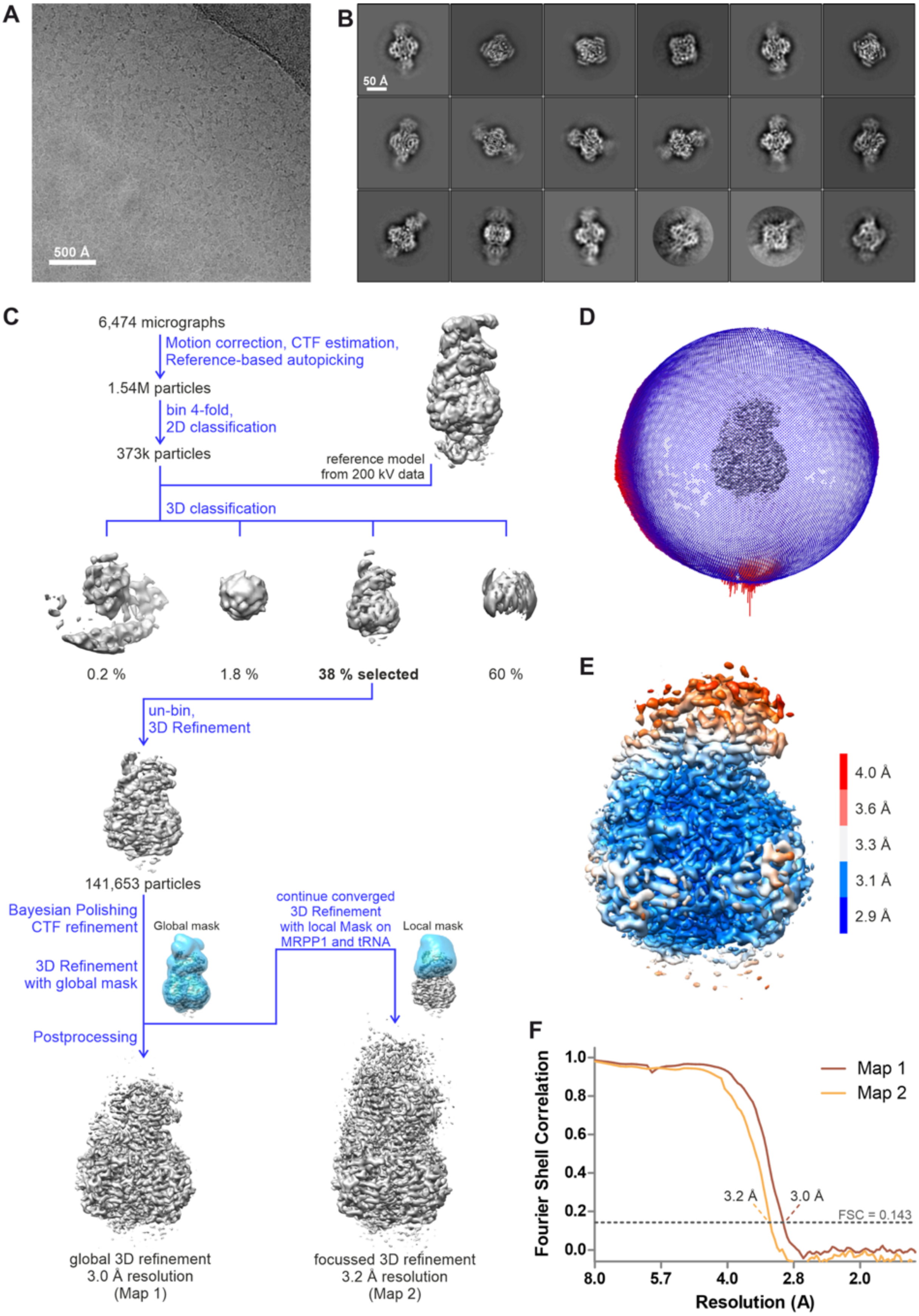
A
500 Å
B
50 Å
C
6,474 micrographs
Motion correction, CTF estimation,
Reference-based autopicking
1.54M particles
bin 4-fold,
2D classification
373k particles
reference model
from 200 kV data
3D classification
0.2 %
1.8 %
38 % selected
60 %
un-bin,
3D Refinement
141,653 particles
Bayesian Polishing
CTF refinement
Global mask
3D Refinement
with global mask
Postprocessing
continue converged
3D Refinement
with local Mask on
MRPP1 and tRNA
Local mask
global 3D refinement
3.0 Å resolution
(Map 1)
focussed 3D refinement
3.2 Å resolution
(Map 2)
D
E
4.0 Å
3.6 Å
3.3 Å
3.1 Å
2.9 Å
F
Fourier Shell Correlation
Resolution (A)
Map 1
Map 2
3.2 Å
3.0 Å
FSC = 0.143
1.0
0.8
0.6
0.4
0.2
0.0
8.0
5.7
4.0
2.8
2.0

**Supplementary Fig. 2: cryo-EM analysis of the TRMT10C/SDR5C1/pre-tRNA$^{Ile}$ complex. a** A micrograph image of the complex. **b** Representative 2D class averages. **c** The data processing work flow. **d** The distribution of orientations of the particles used for the final map, viewed as a surface. **e** Local resolution map of the map. The colour coding scheme indicates the resolution. **f** Fourier shell correlation with and without solvent masks (maps 1 and 2, respectively).

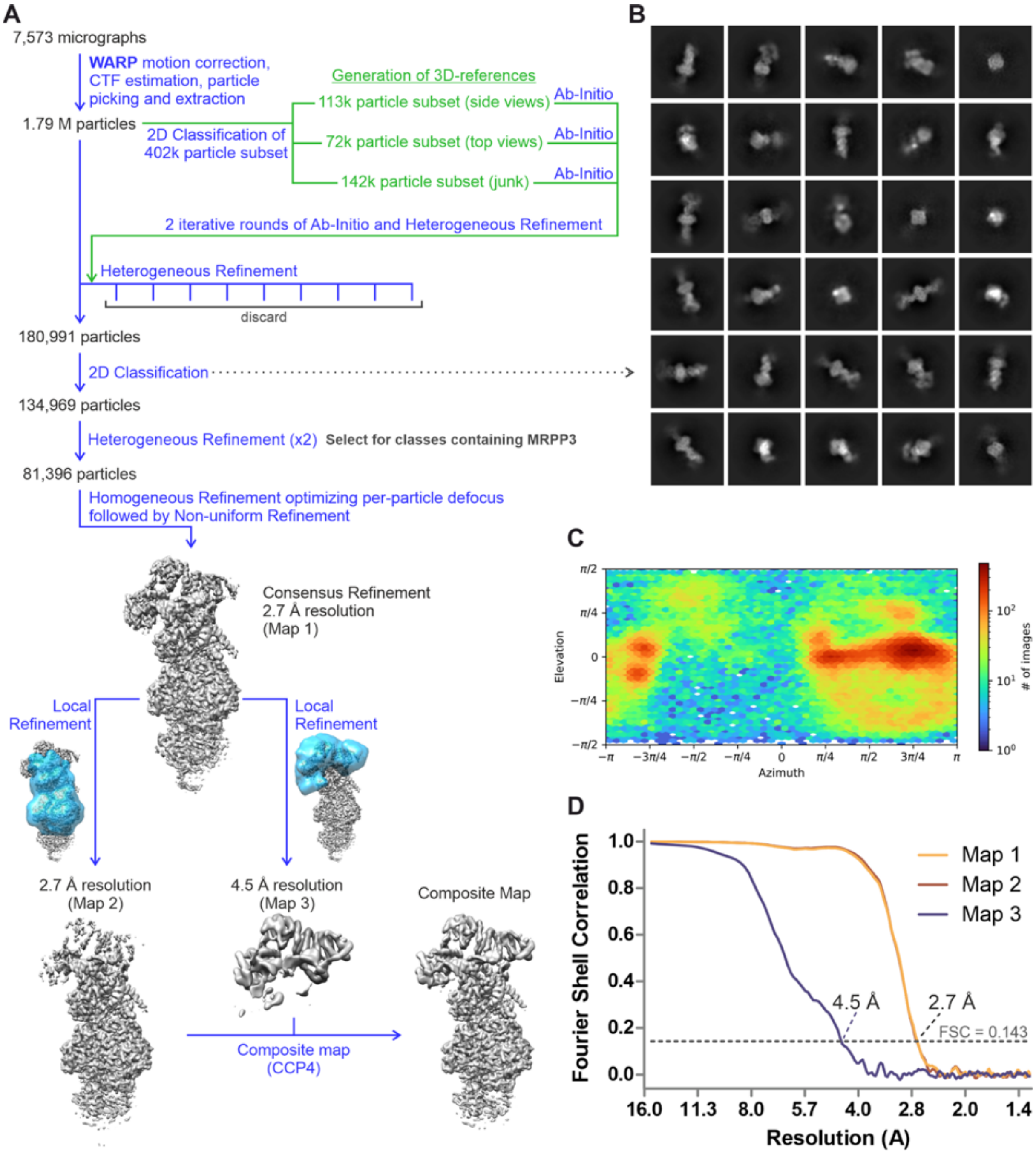

**Supplementary Fig. 3: cryo-EM analysis of the TRMT10C/SDR5C1/pre-tRNA$^{\text{His-Ser}}$/PRORP complex. a** The data processing work flow. **b** Representative 2D class averages. **c** The distribution of orientation of the particles used for the final map in projection. **d** Fourier shell correlation with different particle masks (map 1,2) and without masking (map 3).

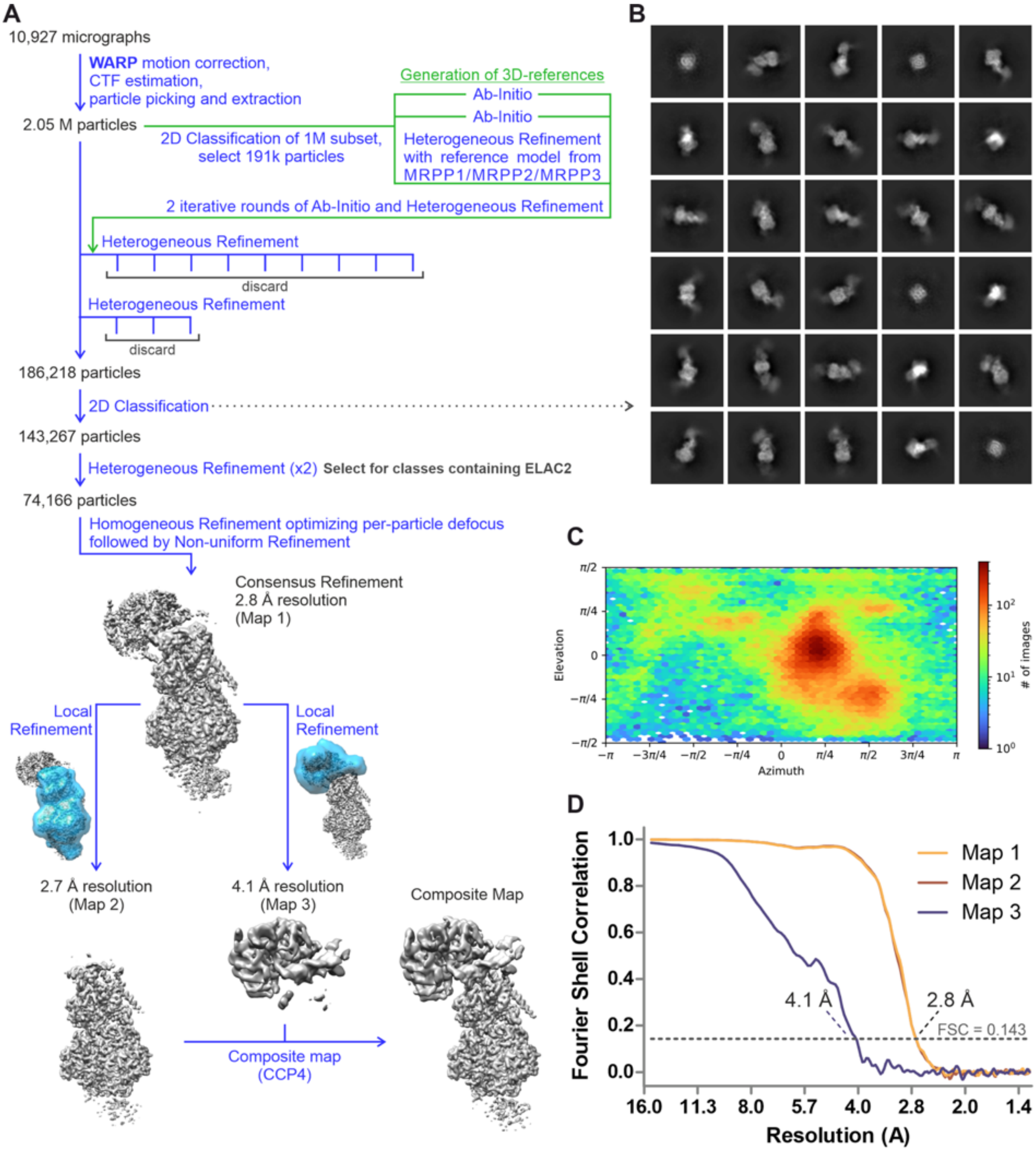

**Supplementary Fig. 4: cryo-EM analysis of the TRMT10C/SDR5C1/pre-tRNA$^{His(0,Ser)}$/ELAC2 complex. a** The data processing work flow. **b** Representative 2D class averages. c) The distribution of orientation of the particles used for the final map in projection. **d** Fourier shell correlation with different particle masks (map 1,2) and without masking (map 3).

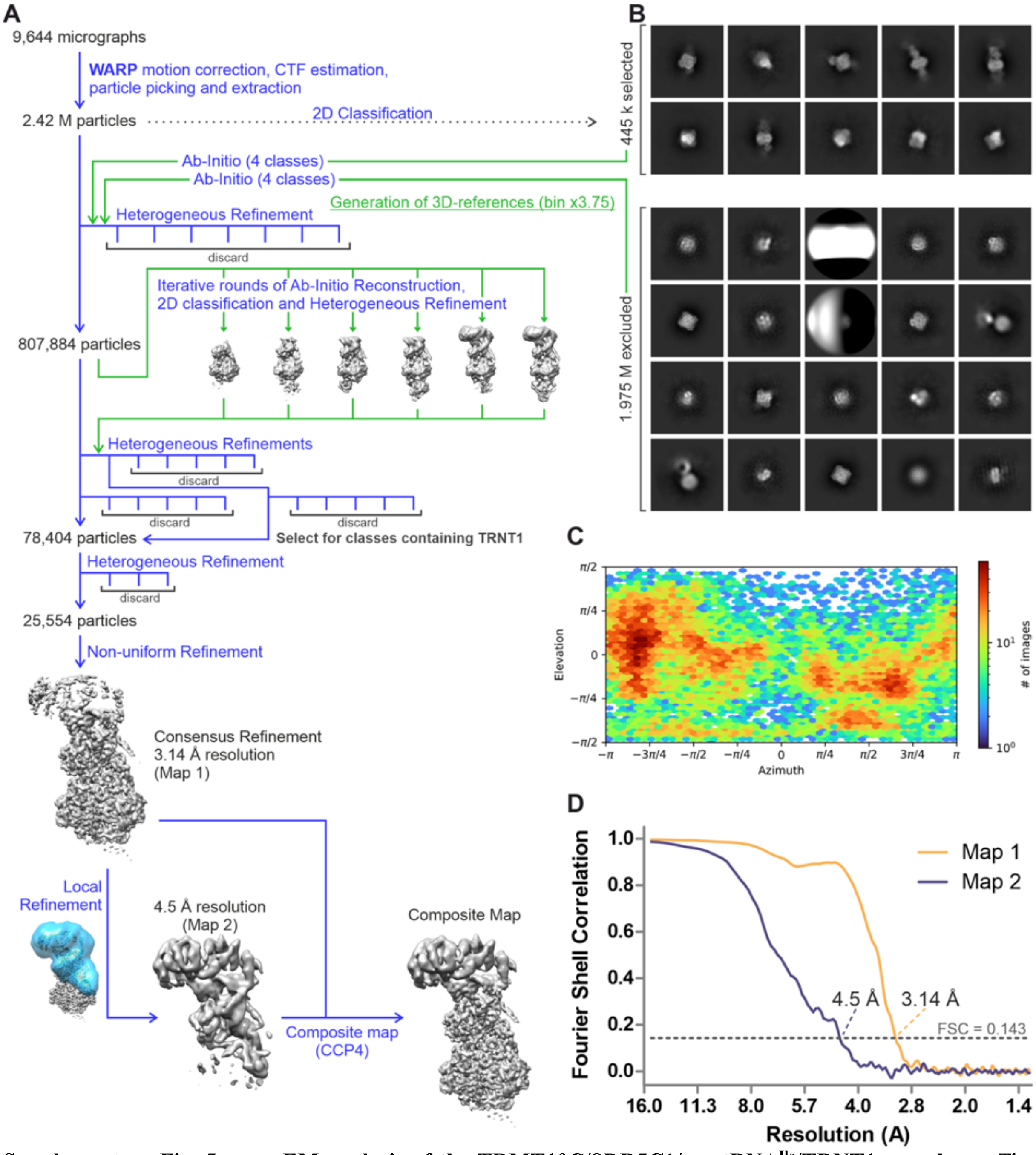

**Supplementary Fig. 5: cryo-EM analysis of the TRMT10C/SDR5C1/pre-tRNA$^{Ile}$/TRNT1 complex. a** The data processing work flow. **b** Representative 2D class averages. **c** The distribution of orientation of the particles used for the final map in projection. **d** Fourier shell correlation with and without particle solvent masks (maps 1,2, respectively).

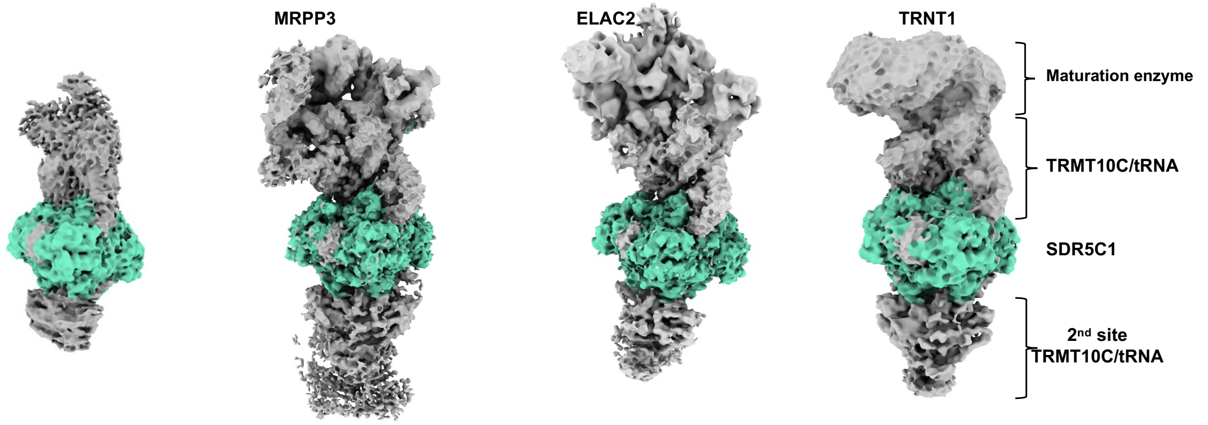

**Supplementary Fig. 6: cryo-EM density maps of the human mt tRNA maturation complexes.** Consensus maps showing the second site of binding of the TRMT10C/tRNA complex on SDR5C1 from left to right: TRMT10C/SDR5C1/pre-tRNA$^{Ile}$, TRMT10C/SDR5C1/pre-tRNA$^{His\text{-}Ser}$/PRORP$_{D479A}$, TRMT10C/SDR5C1/pre-tRNA$^{His(0,Ser)}$/ELAC2$_{H548A}$, TRMT10C/SDR5C1/tRNA$^{Ile}$/TRNT1/CDP.

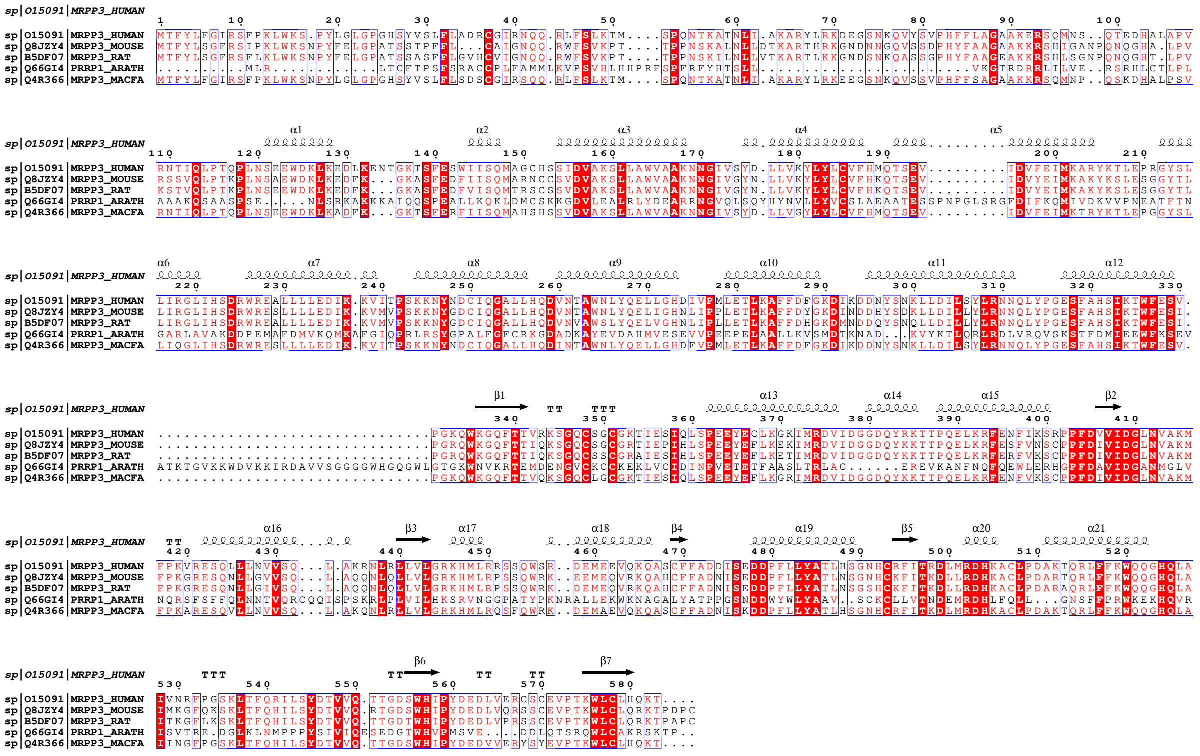

**Supplementary Fig. 7: PRORP sequence conservation.** Multiple sequence alignment of a representative set of PRORP sequences with colour coding from ESPript[59]. The secondary structure of the human PRORP is shown above the alignment and the sequences are numbered according to human PRORP. Abbreviations for sequences are as follows: HUMAN: *Homo Sapiens* (Uniprot O15091), Mouse: *Mus musculus* (Uniprot Q8JZY4), RAT: *Rattus norvegicus* (Uniprot B5DF07), ARATH: *Arabidopsis thaliana* (Uniprot Q66GI4), MACFA: *Macaca fascicularis* (Uniprot Q4R366).

**a**

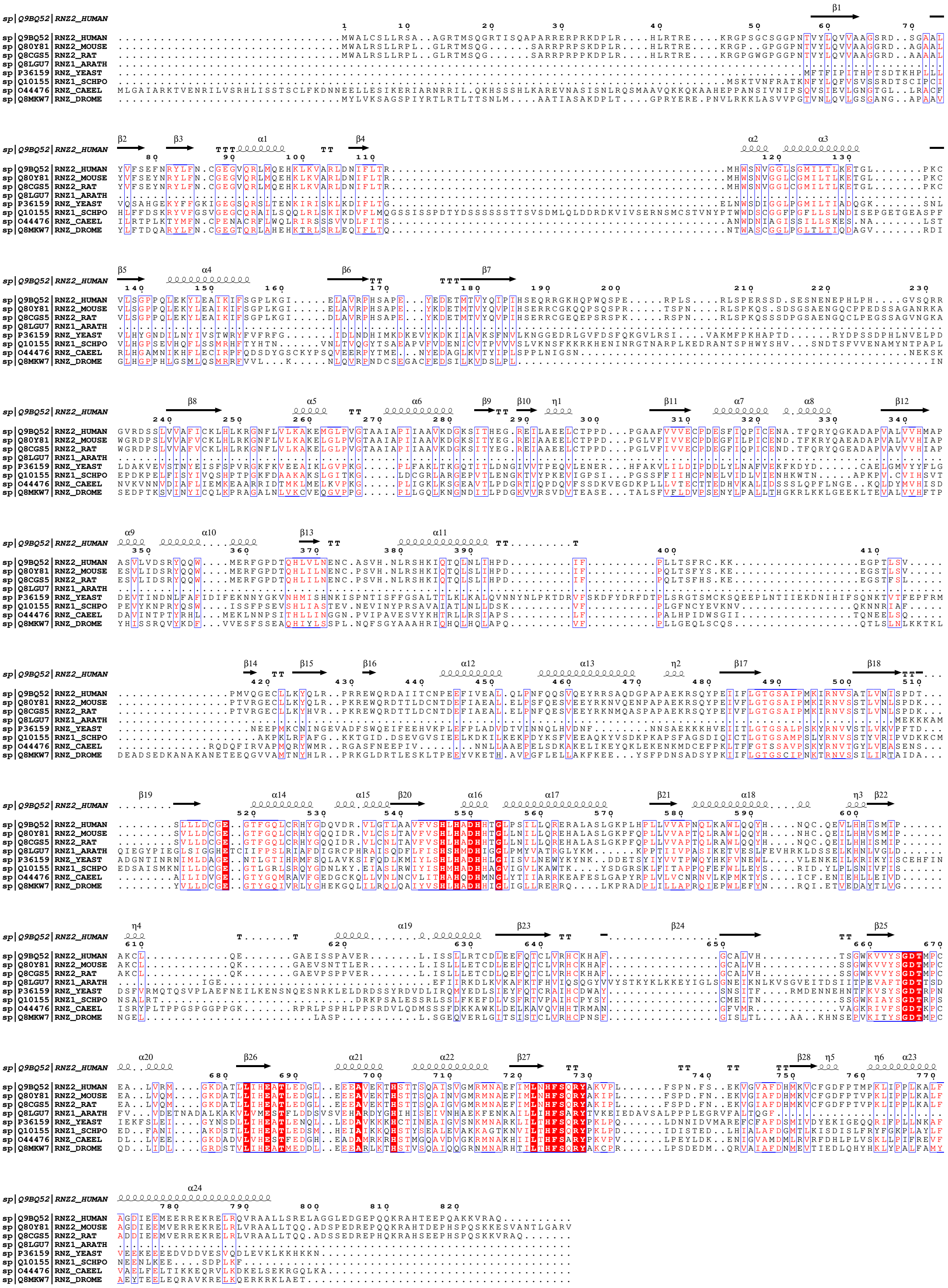

**b**

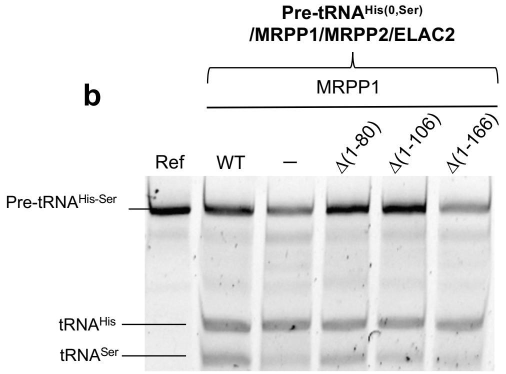

**Supplementary Fig. 8: Conserved residues in ELAC2 and ELAC2 cleavage assays on a pre-tRNA bound in TRMT10C deletion mutants/SDR5C1 complex. a** Multiple sequence alignment of a representative set of ELAC2 sequences with colour coding from ESPript[59]. The secondary structure of the human ELAC2 is shown above the alignment and the sequences are numbered according to ELAC2. Abbreviations for sequences are as follows: HUMAN: *Homo sapiens* (Uniprot Q9BQ52), MOUSE: *Mus musculus* (Uniprot Q80Y81), RAT: *Rattus norvegicus* (Uniprot Q8CGS5), ARATH: *Arabidopsis thaliana* (Uniprot Q8LGU7), YEAST: *Saccharomyces cerevisiae* (Uniprot P36159), SCHPO: *Schizosaccharomyces pombe* (Uniprot Q10155), CAEEL: *Caenorhabditis elegans* (Uniprot O44476) and DROME: *Drosophila melanogaster* (Uniprot Q8MKW7). **b** Cleavage assays of ELAC2 on pre-tRNA$^{His(0,Ser)}$ using TRMT10C deletion mutants in complex with SDR5C1. WT: TRMT10C/SDR5C1/ELAC2/pre-tRNA$^{His(0,Ser)}$, Ref: pre-tRNA$^{His(0,Ser)}$ alone, (-) : ELAC2/pre-tRNA$^{His(0,Ser)}$.

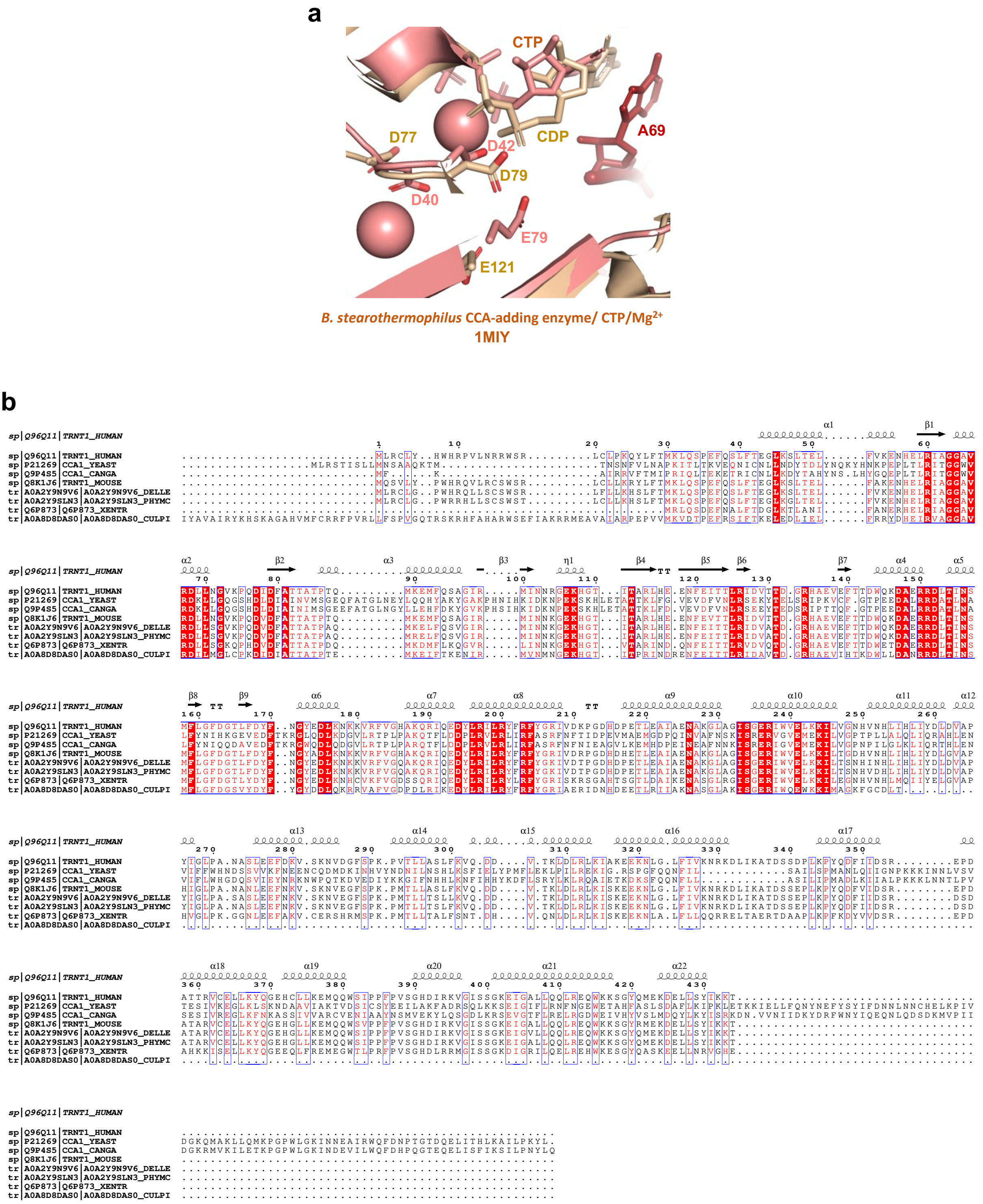

**Supplementary Fig. 9: TRNT1 metal-binding and sequence conservation. a** Superimposition of the TRNT1 active site (in beige) with the crystal structure of *B. stearothermophilus* CCA-adding enzyme in complex with CTP and $Mg^{2+}$ ions (in light magenta, PDB 1MIY). **b** Multiple sequence alignment of a representative set of TRNT1 sequences with colour coding from ESPript[59]. The secondary structure of the human TRNT1 is shown above the alignment and the sequences are numbered according to human TRNT1. Abbreviations for sequences are as follows: HUMAN: *Homo Sapiens* (Uniprot Q96Q11), YEAST: *Saccharomyces cerevisiae* (Uniprot P21269), CANGA: *Candida glabrata* (Uniprot Q9P4S5), MOUSE: *Mus musculus* (Uniprot Q8K1J6), DELLE: *Delphinapterus leucas* (Uniprot A0A2Y9N9V6), PHYMC: *Physeter macrocephalus* (Uniprot A0A2Y9SLN3), XENTR: *Xenopus tropicalis* (Uniprot Q6P873) and CULPI: *Culex pipiens* (Uniprot A0A8D8DAS0).

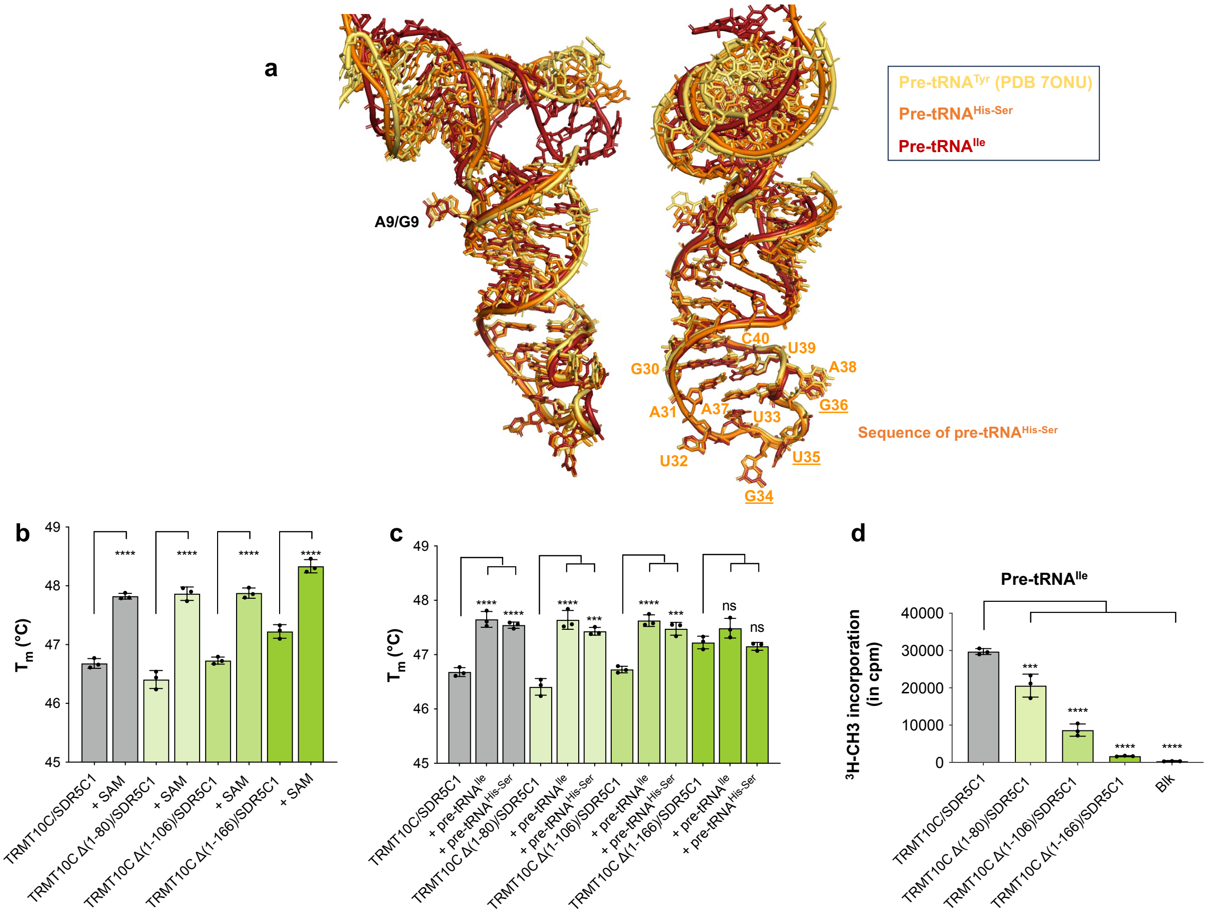

**Supplementary Fig. 10: mt tRNA interactions with TRMT10C.** **a** Superimposition of pre-tRNA$^{His-Ser}$ (in orange), pre-tRNA$^{Ile}$ (in red) and pre-tRNA$^{Tyr}$ (PDB 7ONU, in yellow), tRNA$^{Ser}$ is not shown for the sake of clarity. **b** Melting temperatures (Tm, in °C) of TRMT10C deletion mutants in presence or in absence of SAM, TRMT10C/SDR5C1 WT (in grey), TRMT10C Δ(1-80)/SDR5C1 in light green, TRMT10C Δ (1-106)/SDR5C1 in green and TRMT10C Δ (1-166)/SDR5C1 in lime, measured from DSF experiments. **c** Melting temperatures (Tm, in °C) of TRMT10C deletion mutants in presence of pre-tRNA$^{Ile}$ or pre-tRNA$^{His-Ser}$ measured from DSF experiments, color as in b. **d** MTase assays of TRMT10C deletion mutants on pre-tRNA$^{Ile}$, Blk: assay without pre-tRNA, color as in b. P-values are indicated as follows: ns = P > 0.05, * = P 0.05, *** = P 0.001 and **** = P 0.0001, for n = 3.

**a**

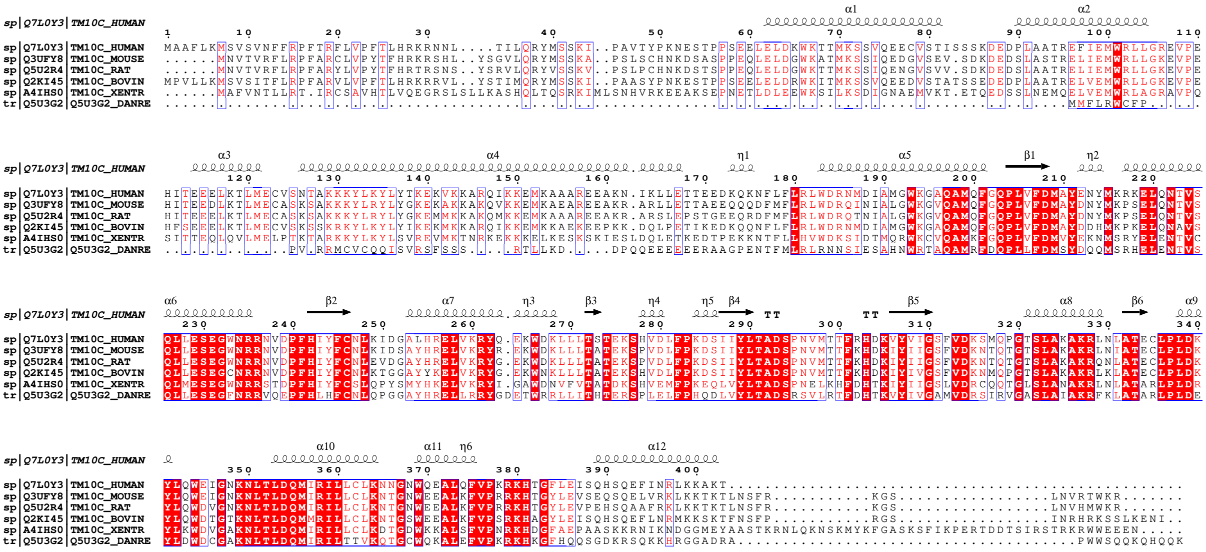

sp|Q7L0Y3|TM10C_HUMAN
sp|Q3UFY8|TM10C_MOUSE
sp|Q5U2R4|TM10C_RAT
sp|Q2KI45|TM10C_BOVIN
sp|A4IHS0|TM10C_XENTR
tr|Q5U3G2|Q5U3G2_DANRE

**b**

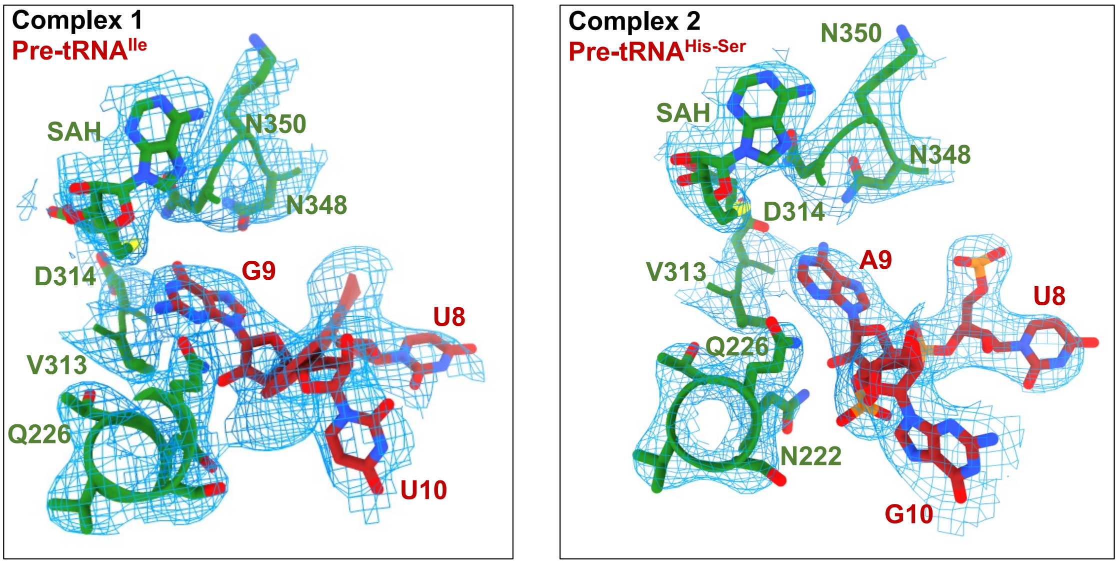

Complex 1
Pre-tRNA^Ile
SAH
N350
N348
D314
G9
U8
V313
Q226
U10
Complex 2
Pre-tRNA^His-Ser
N350
SAH
N348
D314
V313
A9
U8
Q226
N222
G10

**c**

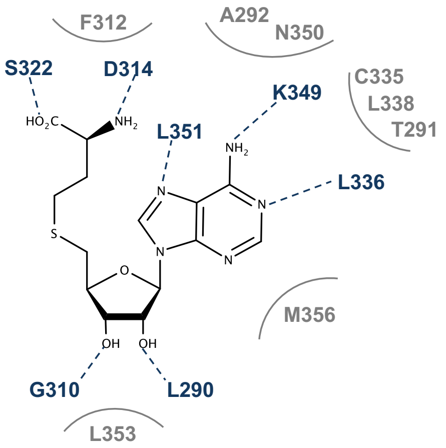

F312
A292
N350
S322
D314
K349
C335
L338
T291
L351
L336
M356
G310
L290
L353

**Supplementary Fig. 11: TRMT10C sequence conservation and active site organization. a** Multiple sequence alignment of a representative set of MRPP1 sequences from eukaryotes with colour coding from ESPript[59]. The secondary structure of the human MRPP1 is shown above the alignment and the sequences are numbered according to human MRPP1. Abbreviations for sequences are as follows: HUMAN: *Homo sapiens* (Q7L0Y3), MOUSE: *Mus musculus* (Uniprot Q3UFY8), RAT: *Rattus norvegicus* (Uniprot Q5U2R4), BOVIN: *Bos taurus* (Uniprot Q2KI45), XENTR: *Xenopus tropicalis* (Uniprot A4IHS0) and DANRE: *Danio rerio* (Uniprot Q5U3G2). **b** Model-to-map fit for G9 and SAH in complex 1 and A9 and SAH in complex 2. **c** Schematic representation of TRMT10C residues interacting with SAH. Dashed lines represent hydrogen bonds between residues and the ligand, residues involved in formation of the binding pocket environment are also shown in grey.

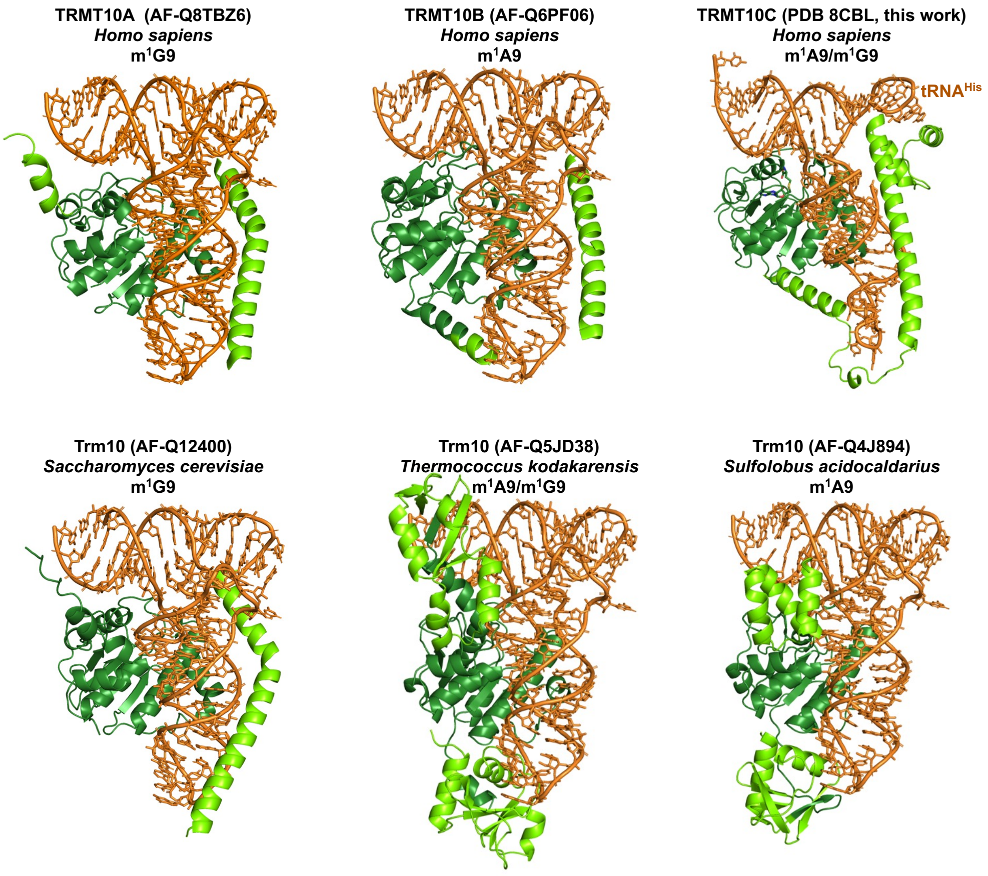

**Supplementary Fig. 12: Models of Trm10-family enzyme/tRNA complexes, highlighting the different strategies to bind tRNA.**

The models were generated using the structure of TRMT10C bound to $tRNA^{His\text{-}Ser}$ (PDB 8CBL, this work), $tRNA^{Phe}$ (PDB 6LVR) and Alphafold models of the Trm10-family enzymes.
